# Exploring Integrative Analysis using the BioMedical Evidence Graph

**DOI:** 10.1101/773911

**Authors:** Adam Struck, Brian Walsh, Alexander Buchanan, Jordan A. Lee, Ryan Spangler, Josh Stuart, Kyle Ellrott

## Abstract

The analysis of cancer biology data involves extremely heterogeneous datasets including information from RNA sequencing, genome-wide copy number, DNA methylation data reporting on epigenomic regulation, somatic mutations from whole-exome or whole-genome analyses, pathology estimates from imaging sections or subtyping, drug response or other treatment outcomes, and various other clinical and phenotypic measurements. Bringing these different resources into a common framework, with a data model that allows for complex relationships as well as dense vectors of features, will unlock integrative analysis. We introduce a graph database and query engine for discovery and analysis of cancer biology, called the BioMedical Evidence Graph (BMEG). The BMEG is unique from other biological data graphs in that sample level molecular information is connected to reference knowledge bases. It combines gene expression and mutation data, with drug response experiments, pathway information databases and literature derived associations. The construction of the BMEG has resulted in a graph containing over 36M vertices and 29M edges. The BMEG system provides a graph query based API to enable analysis, with client code available for Python, Javascript and R, and a server online at bmeg.io. Using this system we have developed several forms of integrated analysis to demonstrate the utility of the system. The BMEG is an evolving resource dedicated to enabling integrative analysis. We have demonstrated queries on the system that illustrate mutation significance analysis, drug response machine learning, patient level knowledge base queries and pathway level analysis. We have compared the resulting graph to other available integrated graph systems, and demonstrated that it is unique in the scale of the graph and the type of data it makes available.

**Highlights:**

- Data resource connected extremely diverse set of cancer data sets
- Graph query engine that can be easily deployed and used on new datasets
- Easily installed python client
- Server online at bmeg.io

**Summary:** The analysis of cancer biology data involves extremely heterogeneous datasets including information. Bringing these different resources into a common framework, with a data model that allows for complex relationships as well as dense vectors of features, will unlock integrative analysis. We introduce a graph database and query engine for discovery and analysis of cancer biology, called the BioMedical Evidence Graph (BMEG). The construction of the BMEG has resulted in a graph containing over 36M vertices and 29M edges. The BMEG system provides a graph query based API to enable analysis, with client code available for Python, Javascript and R, and a server online at bmeg.io. Using this system we have developed several forms of integrated analysis to demonstrate the utility of the system.

## Introduction

Biological data produced by large-scale projects now routinely reaches petabyte levels thanks to major advances in sequencing and imaging. This exponential growth in size is well-documented and is being addressed by multiple big-data initiatives. However, the parallel increase in data heterogeneity is still a major unaddressed issue. With multiple profiling methods, platforms, versions, formats and pipelines, biological data is far from monolithic. The immense and expansive amount of heterogeneous data make it difficult to normalize and integrate data as well as perform integrative analysis across disparate experiments. When faced with these challenges as well as the substantial labor and computation costs, researchers may use only a fraction of publicly available data for their analysis, and will not update their data or analysis as new data becomes available.

Graph databases are useful tools for systems biology analysis where integration of complex data is required^1–3^. In the commercial sector, several major data aggregators have been successfully using graph databases for integration of heterogeneous data. Facebook uses the ‘Social Graph’^4^ to represent the connections between people and their information, while Google’s search engine uses a ‘Knowledge Graph’ to connect various facts about different subjects. This approach is especially powerful when entities in the graph are connected via multiple types of complex, chained interactions. Based on these observations, we have built the BioMedical Evidence Graph (BMEG) to allow for complex integration and analysis of heterogeneous biological data.

The BMEG was created by importing several cancer related resources and transforming them into a coherent graph representation. These resources include patient and sample information, mutations, gene expression, drug response data, genomic annotations and literature based analysis (see Table 1). This Graph contains 15K patients, 54K samples, 4M alleles, 640K drug response experiments and 50K literature derived genotype to phenotype associations.

**Table 1:**
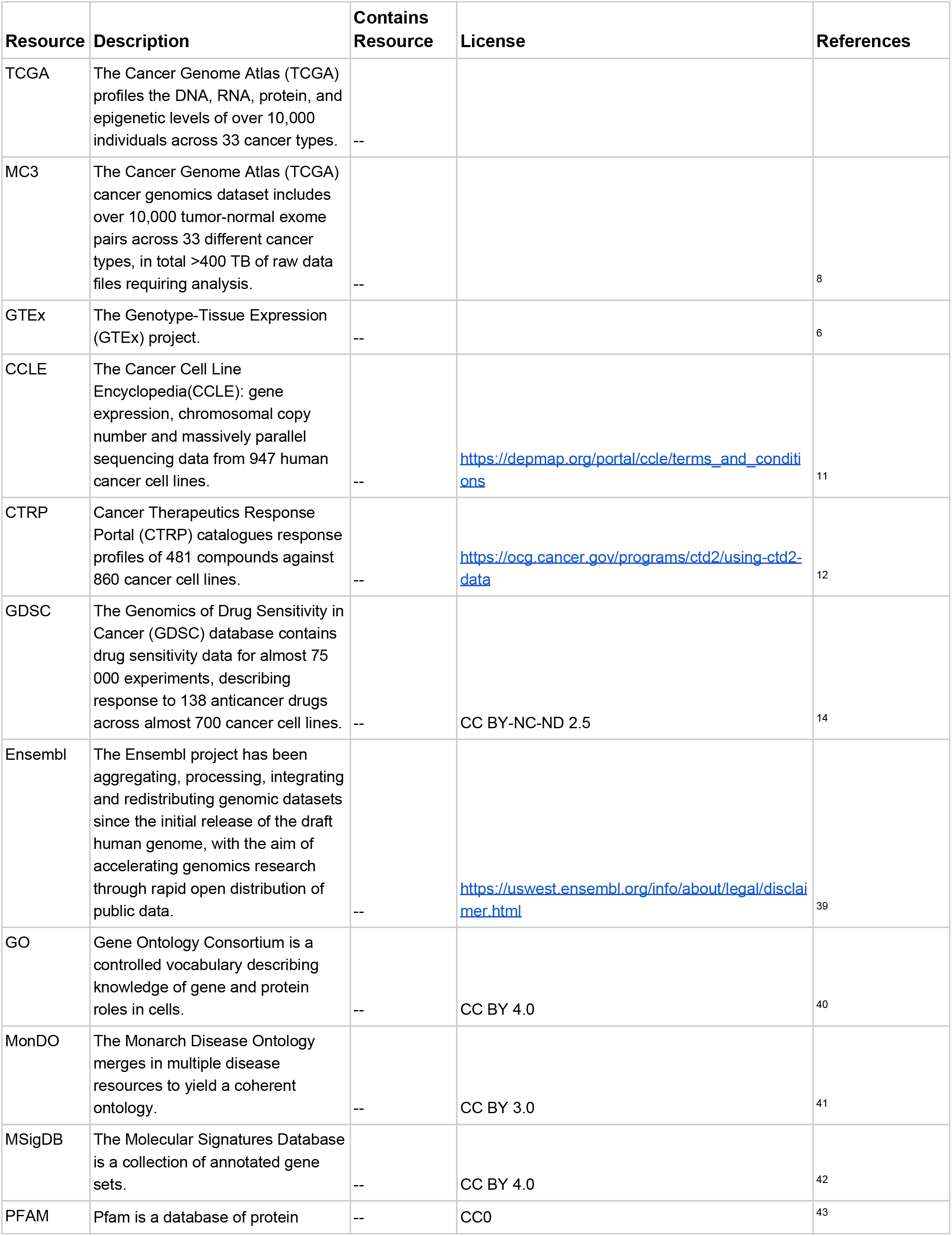
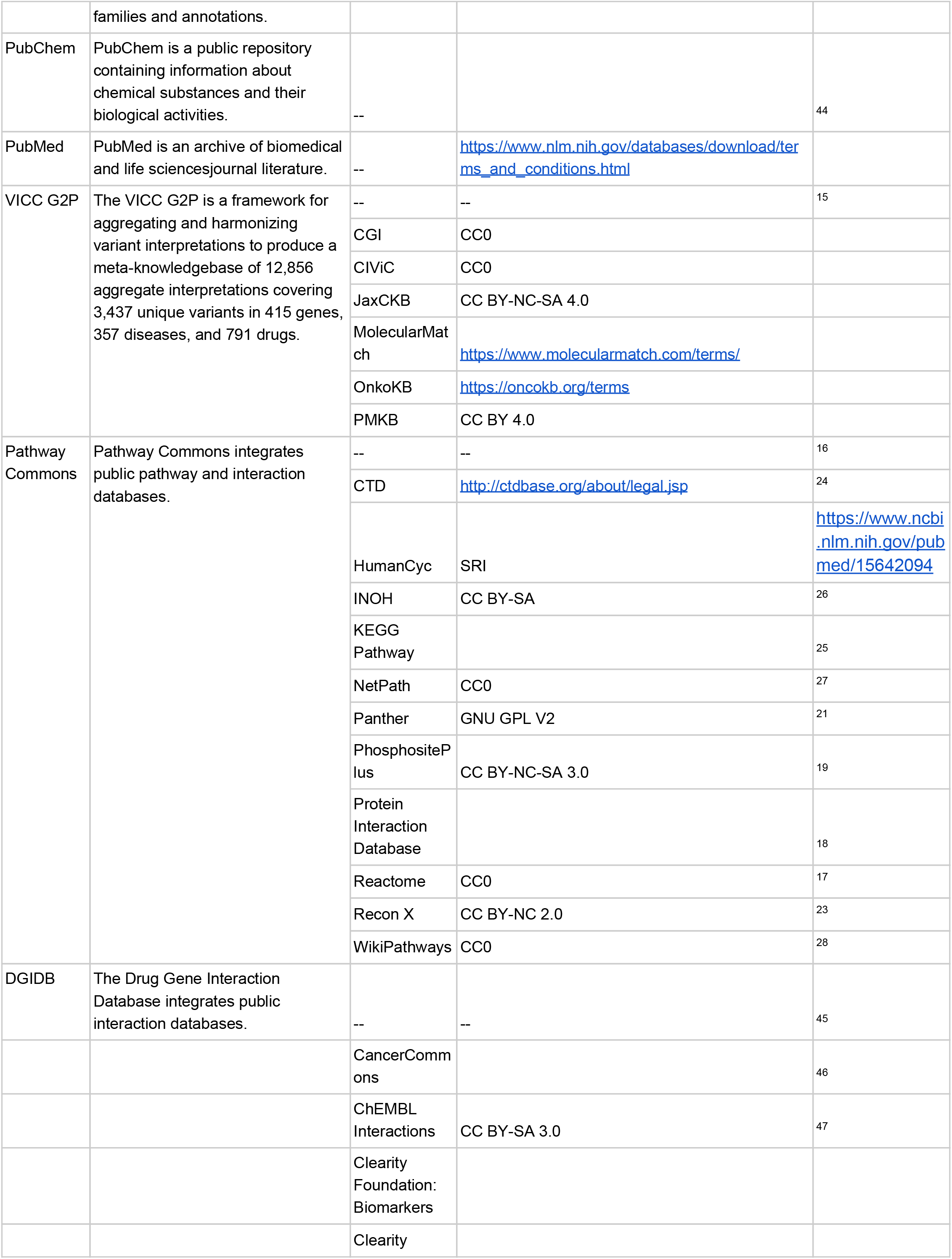
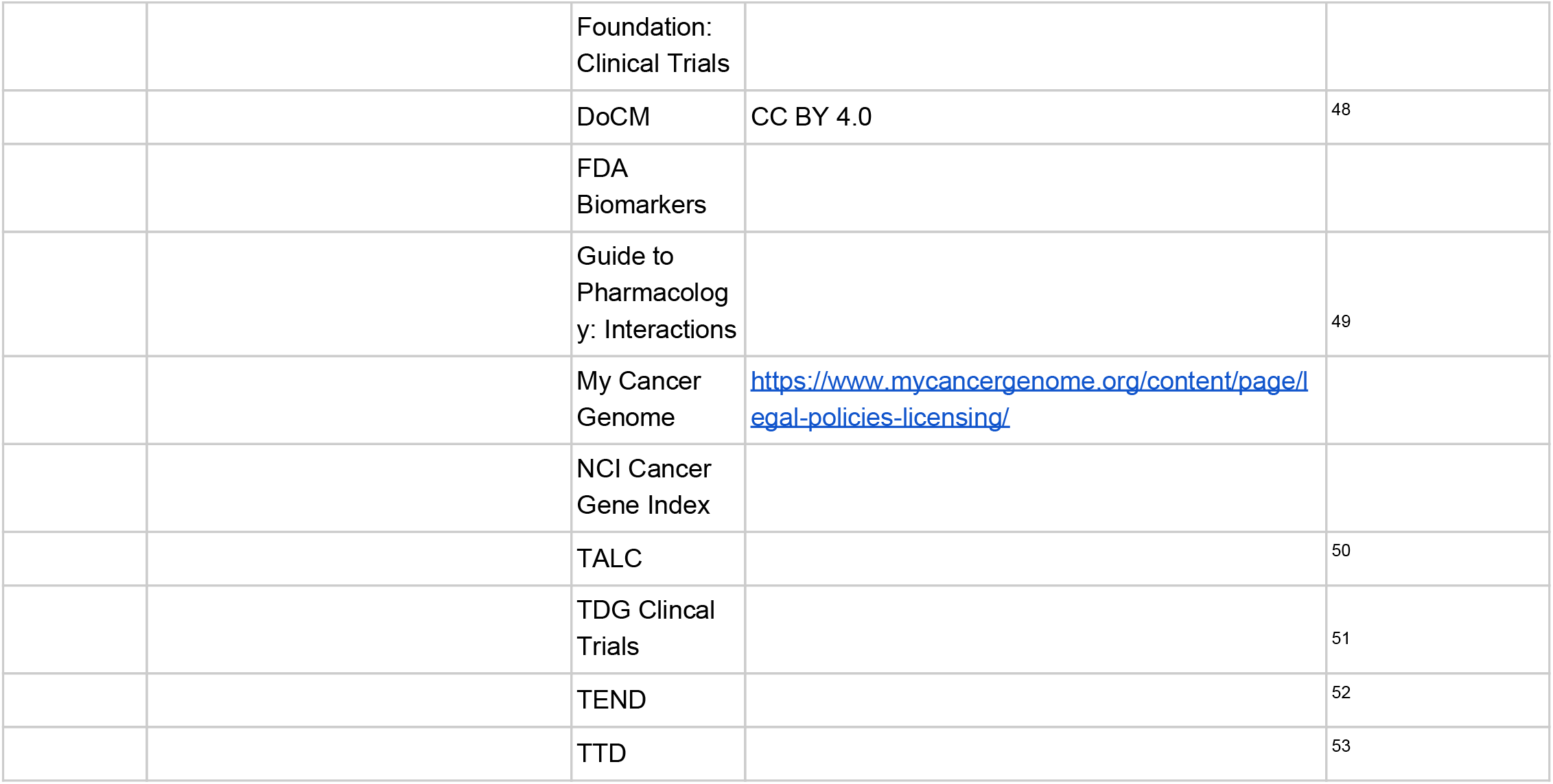
BMEG Data Sources. The different data sources, and their license, that are used to build the BMEG.

To enable analysis and machine learning, our team concentrated on utilizing high quality feature extraction methods applied consistently to all samples. This included identifying the best methods of somatic variant calling and RNA-seq analysis. We utilized open challenges to create leaderboards of the best methods submitted by the community. We then participated in the development of open standards to enable the exchange of genomic associations from cancer knowledge bases.

## Methods

### Graph Schema

Gen3 is a data commons management system developed by the Center for Translational Data Science based on their work for the NCI’s GDC. The BMEG graph schema is described using a JSON schema derived from Gen3 architecture. JSON Schema is a data definition language for describing rules about data structuring. These rules include required fields, data types and field value ranges. The Gen3 system extends JSON schema to add concepts to constructing graph data including database ID alias mapping and edge creation.

At the core of Gen3’s description of TCGA’s metadata, is a tree representing the organization of all the different data elements that make up the program. The tree starts at a top level ‘Program’ node, representing the entire TCGA program, below that are separate projects for each of the different tumor types. Each tumor type is then populated by a number of Cases, which in term have multiple Samples, which can then be subdivided into a number of Aliquots. The BMEG schema builds on this base structure to include data from a number of areas including: 1) Genome Reference, 2) Gene and Pathway Annotations, 3) Somatic Variants, 4) Gene Expression data, 5) Knowledge Bases.

### Data Sources

Initial data sources (see Table 1) for the BMEG were centered on large cohorts of patient-derived samples, with DNA and RNA profiling, cell lines with drug response data and literature-derived drug-phenotype associations. The goal was to provide uniform input data for analysis and machine learning.

### RNA Seq Data

To identify the best methods for RNA analysis, we launched the SMC-RNA challenge, which benchmarked isoform quantification methods to prioritize the methods used for processing data that would be ingested into BMEG. For example, as far as RNA-Seq transcript abundances, we used Kallisto to process the TCGA and CCLE^5^ datasets. Additionally the GTEx project^6^ provided gene-level transcript-per-million mapped reads (TPM) estimates for normal tissues that could be contrasted with tumors. Combinations of these resources provide 36K vertices to the BMEG graph.

### TCGA Metadata

The Genomic Data Commons (GDC) created a data system to track the clinical and administrative meta-data of the TCGA samples and files. We utilized their web API to obtain TCGA patient and sample metadata for the evidence graph.

### TCGA Genomic Data

To determine the best methods for Somatic mutation calling, we partnered with the DREAM consortium, Sage BioNetworks and OICR to launched the ICGC-TCGA Somatic Mutation Calling challenge^7^. Many of the methods evaluated by this effort were incorporated into pipelines that would then be deployed on the TCGA’s 10K exomes as part of the Multi-Center Mutation Calling in Multiple Cancers (MC3) project^8^. The MC3 adds 10K vertices and connects to 3 million alleles (2.6 million distinct) in the graph. For the set of copy number alteration events, we utilized the Gistic2^9^ data from the Broad Institute’s Firehose system.

### Cell Line Drug Response Data

Drug response data has been collated by the DepMap project^10^. This includes response curves, IC50 and EC50 scores from CCLE^11^, CTRP^12, 13^ and GDSC^14^. Additionally, the DepMap meta-data files provided cell line clinical attributes and cross project ID mapping.

### Variant Drug Associations

The Genotype To Phenotype (G2P) schema^15^ was designed to enable a number of different cancer knowledge base resources to be aggregated into a coherent resource. With this resource, the BMEG has aggregated associations from six prominent cancer knowledge bases, including 50K associations vertices.

### Pathway Data

Pathway Commons^16^ aggregates, normalizes and integrates data from 22 public pathway databases. At 1.5 million interactions and 400K detailed biochemical reactions, it is the largest curated pathway databases available. It aggregates pathway relationships from Reactome^17^, NCI Pathway Interaction Database^18^, PhosphoSitePlus^19^, HumanCyc^20^, PANTHER Pathway^21^, MSigDB^22^, Recon X^23^, Comparative Toxicogenomics Database^24^, KEGG Pathway^25^, Integrating Network Objects with Hierarchies^26^, NetPath^27^, and WikiPathways^28^. All of these resources provided 1.9 million vertices to the graph.

### Reference Data

Biological reference data and existing experimental results form a majority of the data stored in the BMEG. These concepts need to be modeled into the graph, and various transformers written to properly translate these concepts. Part of the import pipeline includes Gene, Transcript and Exon annotations, protein and PFAM^29^ assignments as well as Gene Ontology^30^ functional annotations. For this reason, the BMEG standardizes on Ensembl IDs^31^ as the global identifier for various genomic components.

## Graph Databases and Query Languages

To enable various analytical queries, and provide a framework for building new functionality, we developed the GRaph Integration Platform (GRIP) to power queries against the BMEG web resource. GRIP stores multiple forms of data, with the ability to hold thousands of data elements per vertex and per edge of the graph. As applications need, GRIP allows efficient conversion to various data frames for downstream algorithms. This makes the system not only capable of storing sparse relationship data, such as pathways and ontologies, but also dense matrix formatted data, such as gene expression levels for thousands of genes across hundreds of samples.

The query language implements most operations needed for subgraph selection, as well as aggregation features. A general purpose endpoint places more emphasis on the client side building smart queries to obtain the data they need, rather than having custom server side components provide specialized facets to the users. Because of this, clients can easily create new queries, unanticipated by the server developers, and have them still work. We have made the API available via Python, Javascript and R clients. GRIP is written in the GO programming language, and compiles to a single static binary, which means that it can be installed onto a system with little to no dependencies.

## Results

Every different example use case demonstrates the utility of the BMEG dataset and its query engine. GripQL is a traversal based graph selection language based on Gremlin^32^. The user describes a series of steps that will be undertaken by a ‘traveler’. An example traversal would start on a vertex with label ‘project’, go out to edges labeled ‘samples’, then go out along edges labeled ‘aliquots’. The engine then scans the graph for all valid paths that can be completed given the instructions. Each of the traversal descriptions is based on the graph schema seen in Figure 1. The commands are written using the Python version of the client, but could be executed similarly in R or Javascript. The API provides a ***getSchema*** method which describes the different types of vertices, their properties and the edges that connect them.

**Figure 1:**
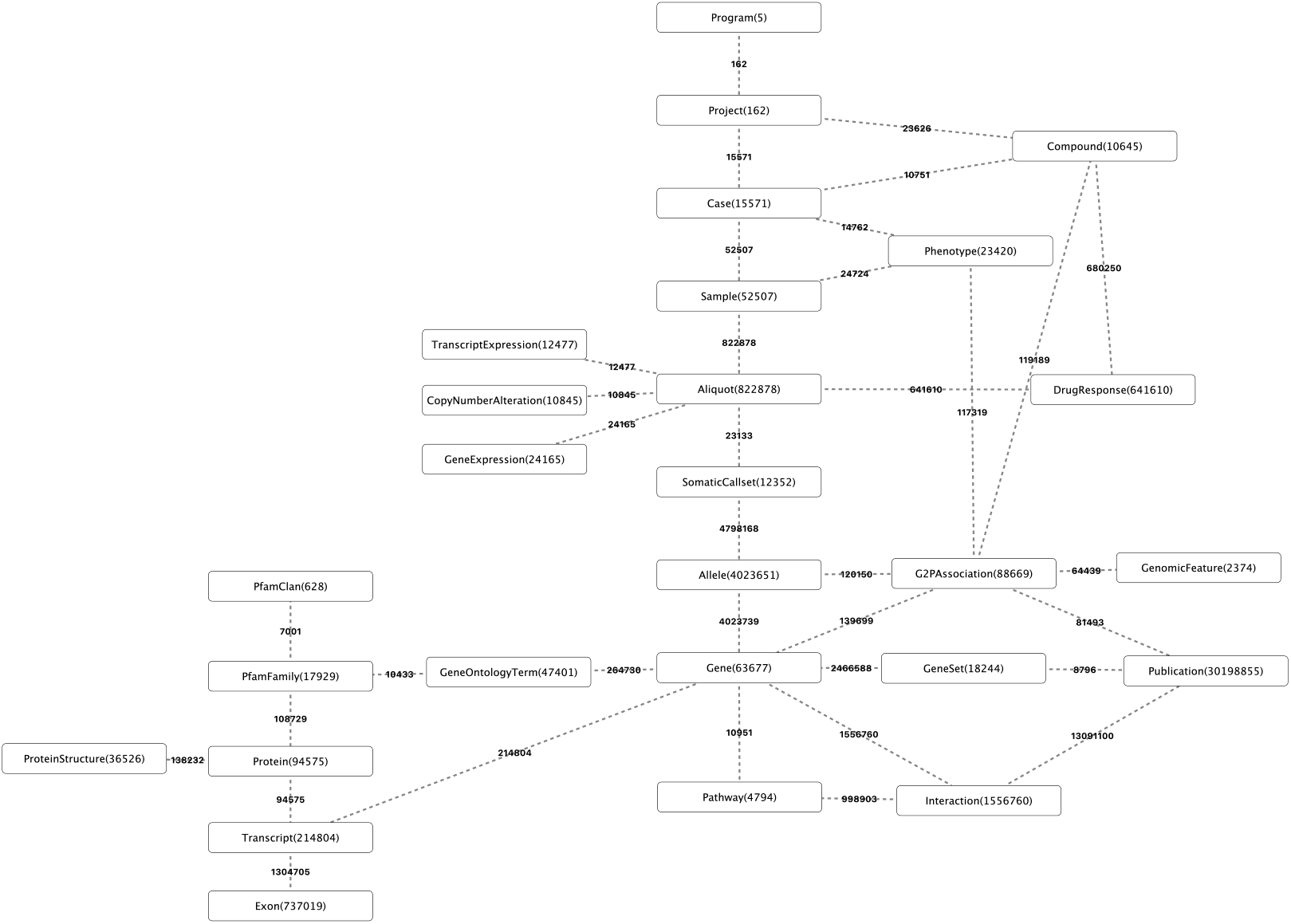
Graph Schema. The vertex types and connections of the BMEG. The numbers represent the count for each vertex type, and the counts on the edges represent the total number of connections between those two vertex types.

**Figure 2:**
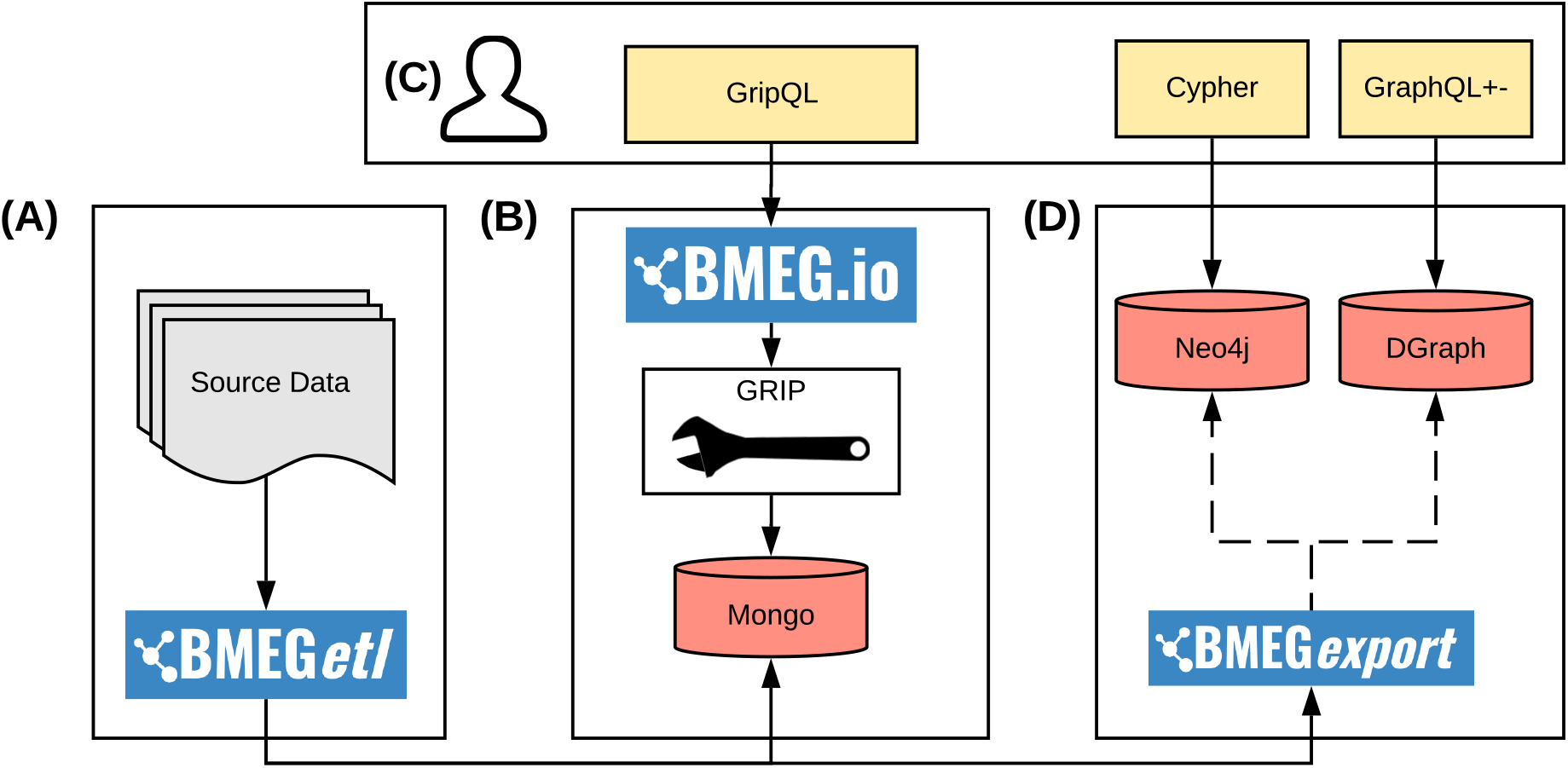
BMEG Architecture. High Level architecture diagram of the BMEG. A) represents the Extract Transform Load (ETL) processes that are used to build the graph, B) the database and query engine used to power the bmeg.io site, C) the different client site options for communicating with the system, D) Other graph engines that can be used with the BMEGexport code to move the BMEG data to other graph databases.

**Figure 3:**
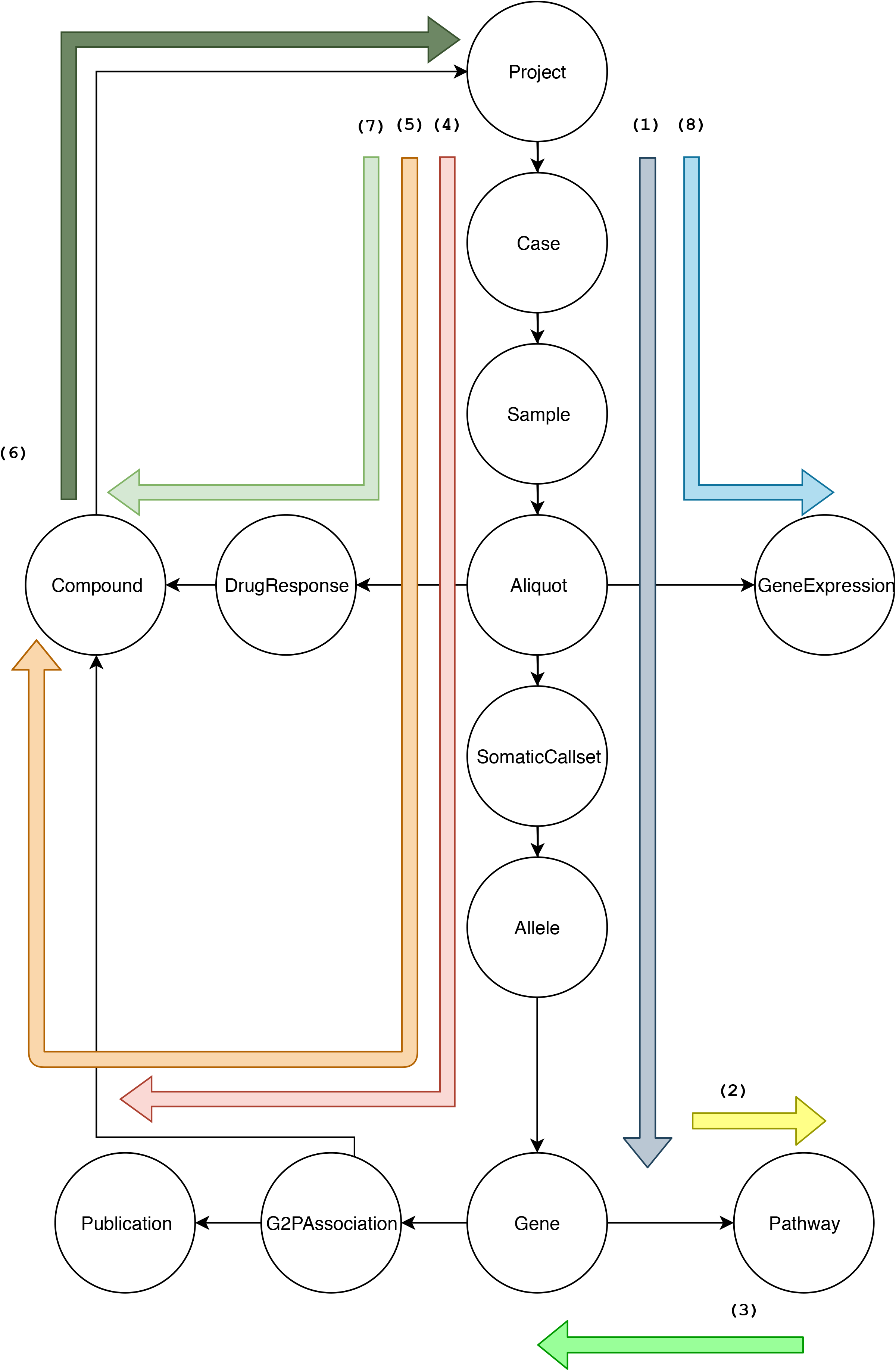
Example Queries. A diagram showing how each of the different queries described in this paper traverse the graph. Each separate query is labeled by the equation number in the text.

Our analysis begins with counting the mutations per gene in a cancer cohort. As seen in (1), this query starts on a node of a project, in this case *TCGA-BRCA*, and then follows the path to the cases that belong to the project, then to the samples and finally to the aliquots. As it passes the *Sample* node, it filters for tumor samples. Once on the aliquot node, it continues to the *SomaticCallset*, which represents sets of variants produced by a single mutation calling analysis. The traversal then identifies the edges that connect the *SomaticCallet* to different alleles, this time using the ***outE*** command to land on the edge, rather than the destination vertex. With the gene ID in hand, we then uses the ***aggregate*** method to count up the various terms that occur in the *enembl_gene* field.

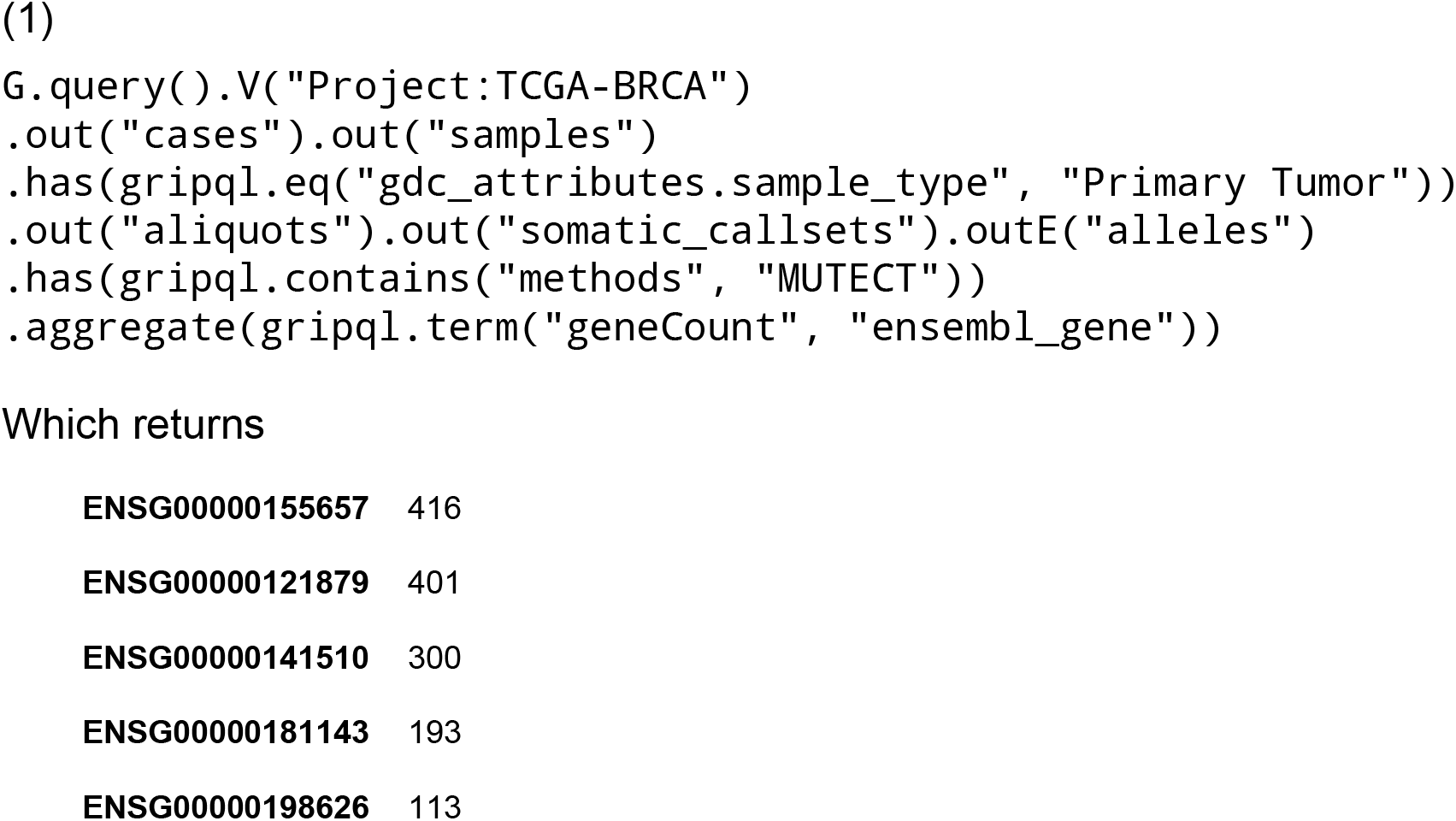

listing the number of variant alleles found for each gene.

We can then inspect the most frequently impacted pathways. First by identifying which pathways each of the mutated genes belong to in (2) and then by normalizing by the number of genes per pathway in (3). To identify the pathways involved in mutations, we provide a list of all mutated genes, find their associated pathways and retrieve the tuples of every gene-pathway pair, using the ***as_*** command (the underscore is added to avoid clashing with the python reserved word ***as***) to store the gene and then using the ***render*** function to display only the data we require.

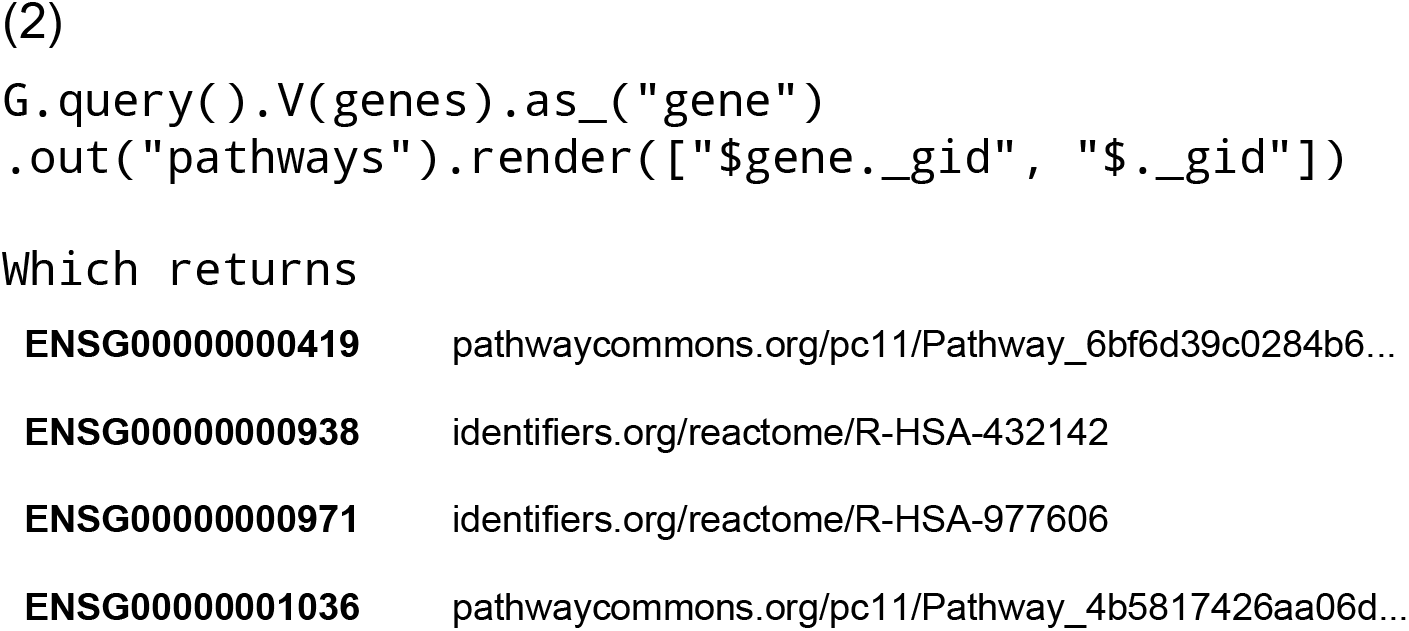

listing all of the pathways for which each gene is a member.

This information can be combined with the previous table to calculate the mutations per pathway. To normalize these values we count the number of genes per pathway. The traversal first starts on the *Pathway* vertex marks for later retrieval using the ***as_*** command. Once the travel has split and moved out to the multiple child *Gene* vertices, the ***select*** command recalls the stored pathway vertex and moves the traveler back. As this point, an ***aggregation*** is called to count the number of travelers on each *Pathway* vertex.

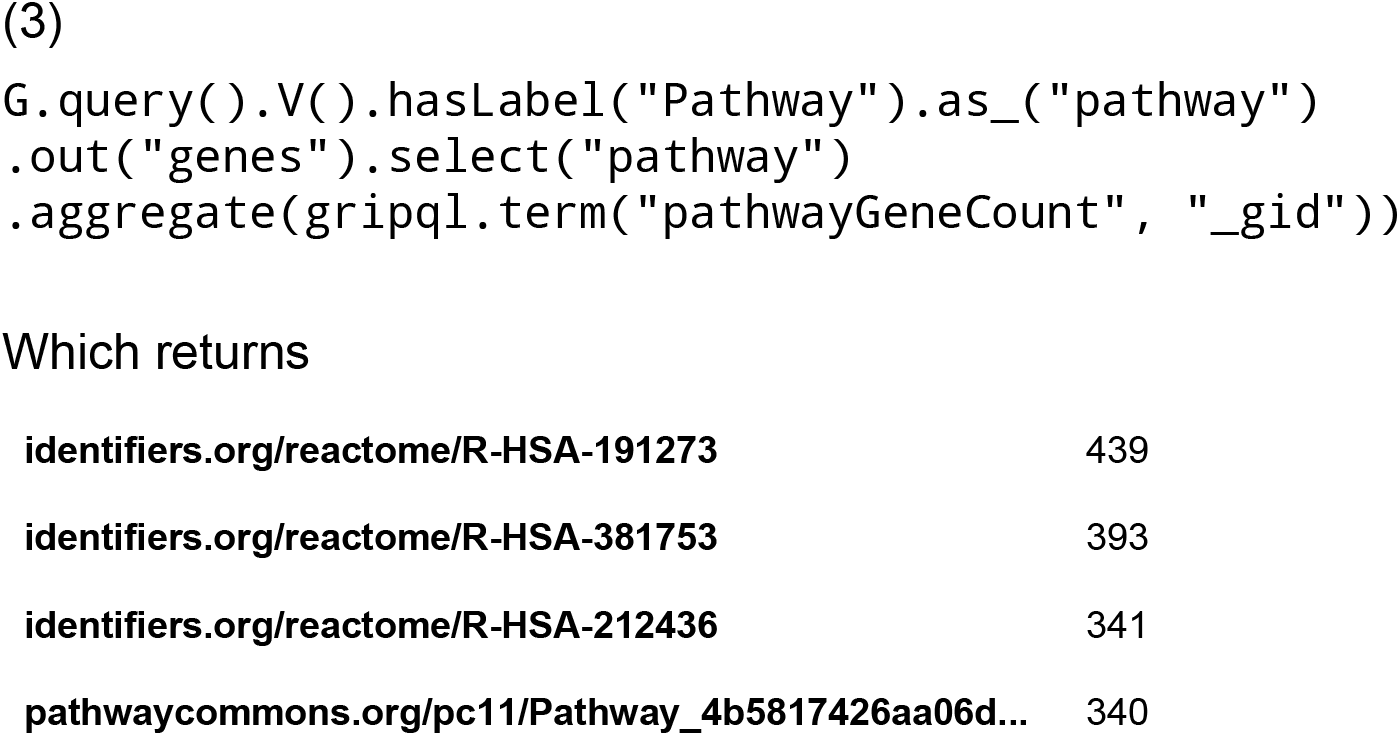

listing the number of mutations for each pathway found.

We can also connect the sets of mutations in the BRCA cohort to the most connected papers, as linked by the G2P associations in (4). In this use case, the aggregate method is called on the special ***_gid*** variable which represents a unique Global ID for each vertex.

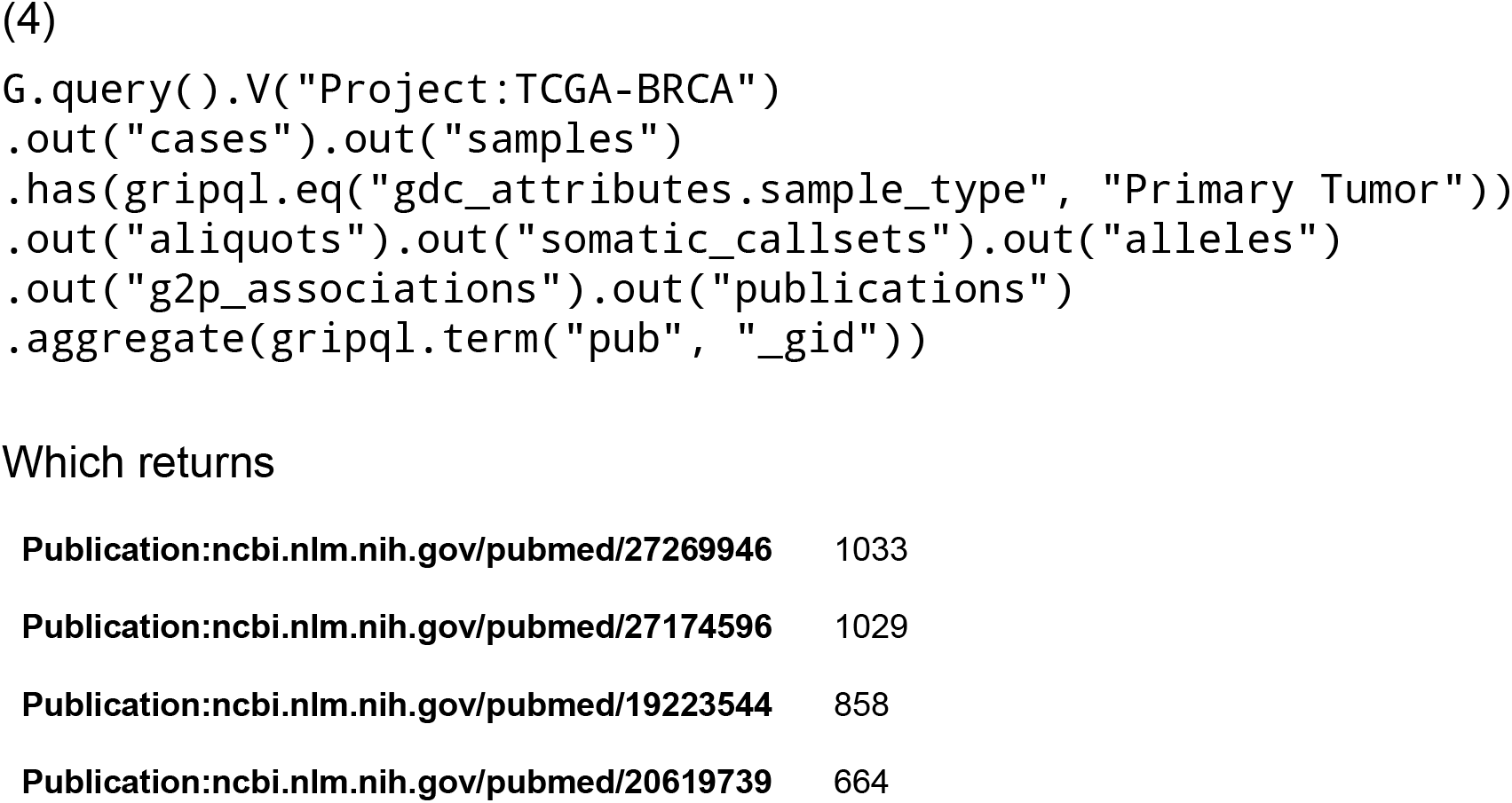

That lists the number of mutations for all genes connected in each of the returned papers.

The G2P associations also connect to compounds that are linked to phenotypes based on specific alleles in (5). This traversal is much like (4), however it also includes a ***distinct*** operation to identify unique pairs of cases and compounds. If there are multiple known association records from different publications and these publications link one allele to the same drug response phenotype, then only one relationship will be noted per patient.

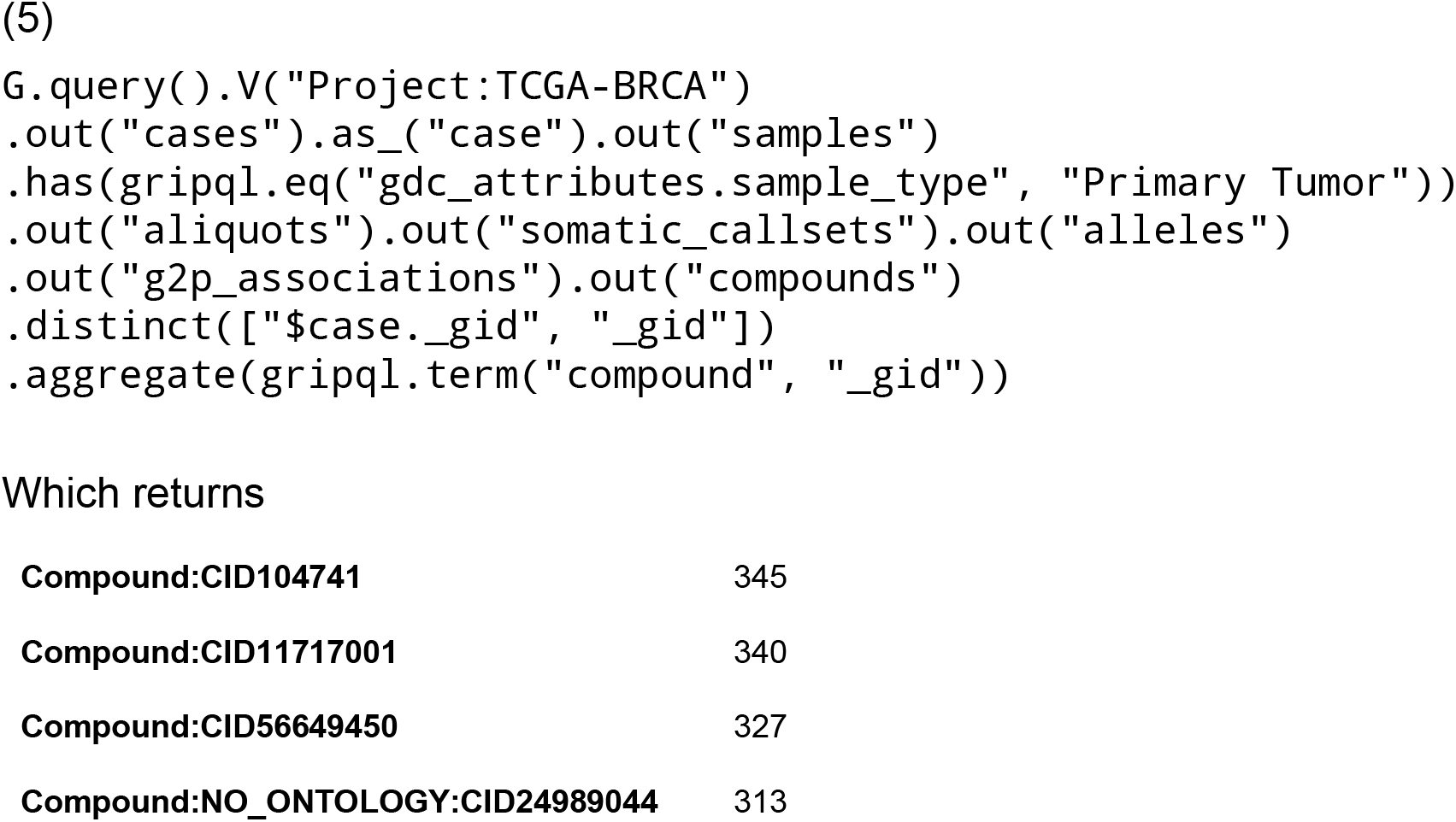

listing the number of times compounds where associated to mutations in patients.

We can now take the most commonly linked drugs, in a list named *compounds*, that we found in (5) and identify the ones that have also been part of cell line drug response testing in (6). We limit our query to cell lines that were studied as part of the CTRP set of breast cancer cell lines.

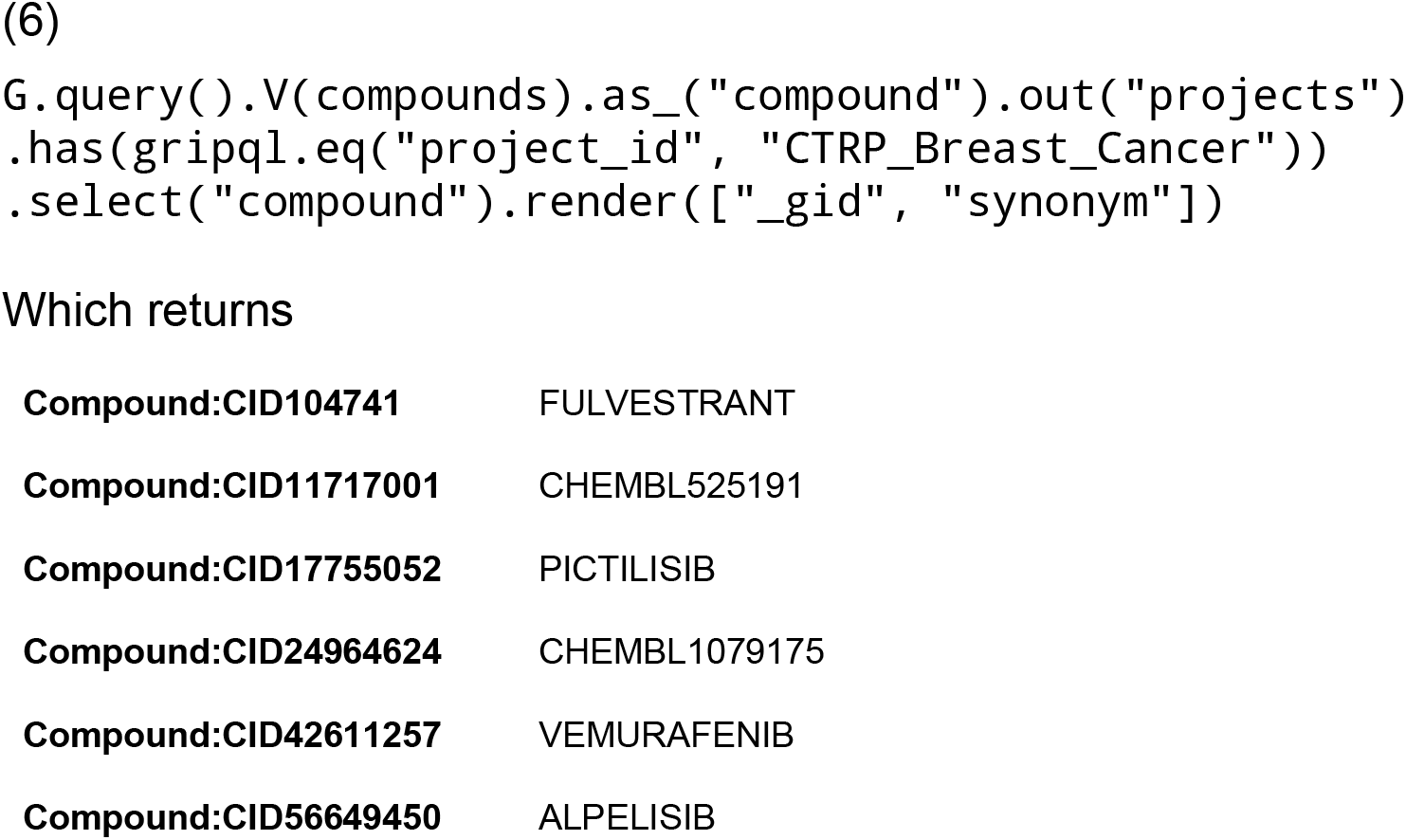

listing those drugs that were profiled in the CTRP effort.

At this point, we find that Compound:CID104741 (Fulvestrant) was on our list of referenced drugs and was studied as part of CTRP. In (7) we look for the EC50 values for the samples in the breast cancer cell lines that were tested against Fulvestrant. This also includes a call to the ***render*** method which shapes the output into a custom JSON structure. In this case, it forms a tuple with the stored sample ID and EC50 value. The listed of tuples returned by the client can then be passed directly into a Pandas DataFrame^33^.

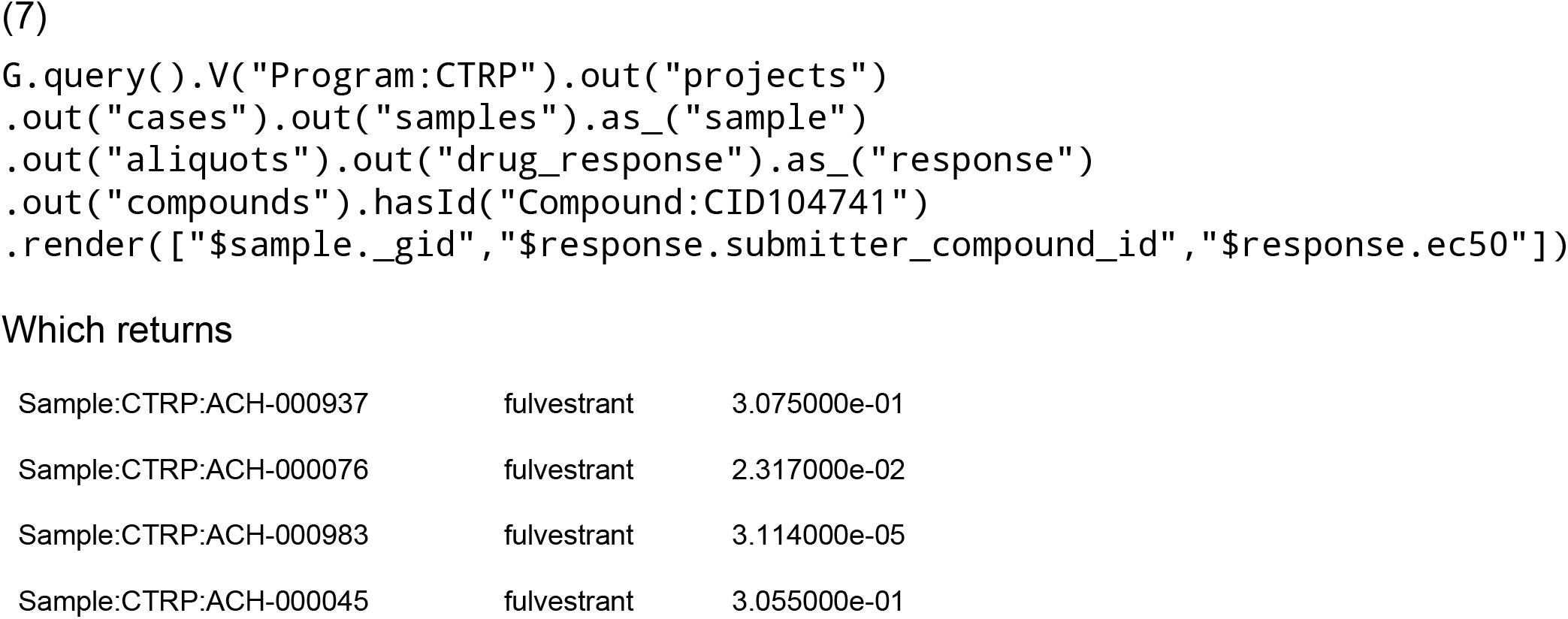

listing the EC50 values for each of the BRCA cell lines to the Fulvestrant agent.

With the drug response values in hand, we can then look for associated transcriptomic data. There is no direct RNA sequencing available from the CTRP project, however many of the cell lines used in the CTRP project were assayed as part of the CCLE project. To identify these samples, in (8) we follow the edge connecting the list named *samples* found in (7) to their parent cases. We then follow the *same_as* edge to identify *Case* vertices in other projects that are the same as the ones we started on, and then follow the tree down to the *GeneExpression* node to obtain the expression values and then link them to our sample IDs. Again, we can use the render function to return properly formatted data structures that can be passed directly into Pandas.

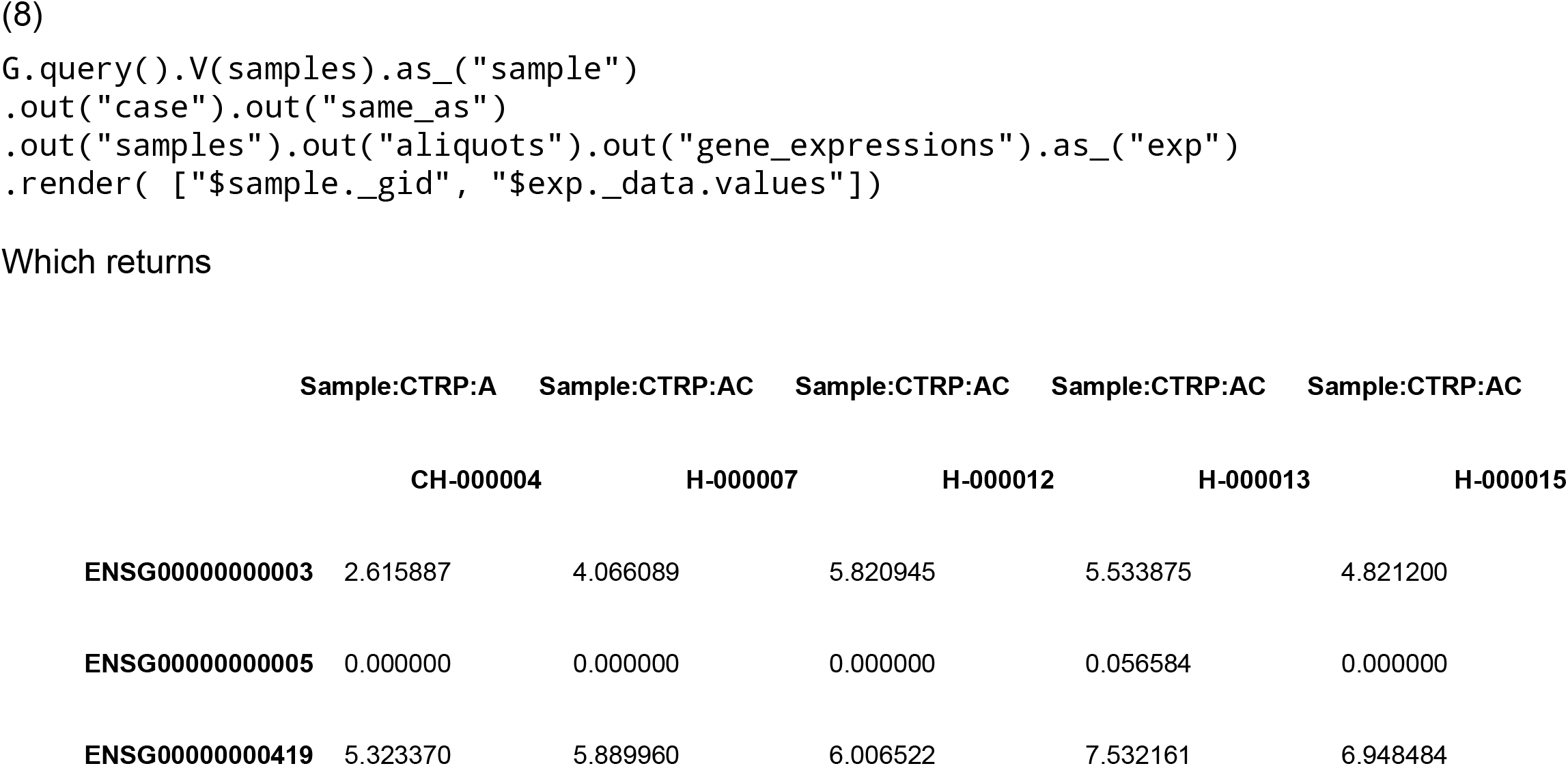

listing the expression values of each gene across cell lines with variants in CTRP and RNA in CCLE. The resulting matrix can be used to develop transcriptome based drug response models.

### Data Releases

The BMEG resource was designed with portability and openness in mind. The graph query engine that runs the system is open source and easy to install, while all of the compiled source files are made available for bulk download. This allows other researchers to build on our existing system, and reuse the data we’ve collected. Because graph data can be represented by a number of different query engines, we also developed translations of the BMEG resource. Part of the BMEG toolkit is a set of scripts to translate the data set and load it into other graph database systems including Neo4J and Dgraph.

## Discussion

Recently, a number of graph-based data integration projects have appeared, including biograkn^34^, Biograph^35^, Bio4j^36^, Bio2RDF^37^, Hetionet^38^. Many of these systems were built to aggregate pathway and genotype/phenotype linkages. BMEG is unique from these efforts, in that its primary use case is to drive analysis and machine-learning from actual samples. The BMEG holds genomic, transcriptomic and phenotypic data from cancer cases as well as cell line samples. This data is meant to provide a starting point for discovery, and generation of new models, rather than simply a repository of existing models. The core of the BMEG idea is to define a coherent input data set to enable various downstream analysis possibilities.

With the primary layer of data in place, the next step is to enable online machine learning methods and do a comparative analysis of the patterns learned. The next steps, which are currently being developed, will see machine learning based predictions join the graph. These derived associations will connect samples to phenotypes similar to the way that the G2P edges connect samples to drug sensitivity. With these novel phenotypic annotations available, we will be able to observe predicted trends across cohorts and identify new patterns.

## Supporting information

IPython Examples

## Author Contributions

Conception and design: Josh Stuart, Kyle Ellrott Collection and assembly of data: Adam Struck, Brian Walsh, Alexander Buchanan, Ryan Spangler Data analysis and interpretation: Adam Struck, Brian Walsh, Jordan Lee Manuscript writing: All authors Final approval of manuscript: All authors Accountable for all aspects of the work: All authors

## Bibliography

1. Yoon B-H, Kim S-K, Kim S-Y: Use of Graph Database for the Integration of Heterogeneous Biological Data. Genomics Inform 15:19–27, 2017

2. Have CT, Jensen LJ: Are graph databases ready for bioinformatics? Bioinformatics 29:3107–3108, 2013

3. Lysenko A, Roznovăţ IA, Saqi M, et al: Representing and querying disease networks using graph databases. BioData Min 9:23, 2016

4. Ugander J, Karrer B, Backstrom L, et al: The Anatomy of the Facebook Social Graph [Internet]. arXiv [csSI], 2011Available from: http://arxiv.org/abs/1111.4503

5. Tatlow PJ, Piccolo SR: A cloud-based workflow to quantify transcript-expression levels in public cancer compendia. Sci Rep 6:39259, 2016

6. GTEx Consortium: The Genotype-Tissue Expression (GTEx) project. Nat Genet 45:580–585, 2013

7. Boutros PC, Ewing AD, Ellrott K, et al: Global optimization of somatic variant identification in cancer genomes with a global community challenge. Nat Genet 46:318–319, 2014

8. Ellrott K, Bailey MH, Saksena G, et al: Scalable Open Science Approach for Mutation Calling of Tumor Exomes Using Multiple Genomic Pipelines. Cell Syst 6:271–281.e7, 2018

9. Mermel CH, Schumacher SE, Hill B, et al: GISTIC2.0 facilitates sensitive and confident localization of the targets of focal somatic copy-number alteration in human cancers. Genome Biol 12:R41, 2011

10. Ghandi M, Huang FW, Jané-Valbuena J, et al: Next-generation characterization of the Cancer Cell Line Encyclopedia. Nature 569:503–508, 2019

11. Barretina J, Caponigro G, Stransky N, et al: The Cancer Cell Line Encyclopedia enables predictive modelling of anticancer drug sensitivity. Nature 483:603–607, 2012

12. Basu A, Bodycombe NE, Cheah JH, et al: An interactive resource to identify cancer genetic and lineage dependencies targeted by small molecules. Cell 154:1151–1161, 2013

13. Rees MG, Seashore-Ludlow B, Cheah JH, et al: Correlating chemical sensitivity and basal gene expression reveals mechanism of action. Nat Chem Biol 12:109–116, 2016

14. Yang W, Soares J, Greninger P, et al: Genomics of Drug Sensitivity in Cancer (GDSC): a resource for therapeutic biomarker discovery in cancer cells. Nucleic Acids Res 41:D955–61, 2013

15. Wagner AH, Walsh B, Mayfield G, et al: A harmonized meta-knowledgebase of clinical interpretations of cancer genomic variants [Internet]. bioRxiv 366856, 2018[cited 2019 Sep 13] Available from: https://www.biorxiv.org/content/10.1101/366856v2

16. Cerami EG, Gross BE, Demir E, et al: Pathway Commons, a web resource for biological pathway data. Nucleic Acids Res 39:D685–90, 2011

17. Fabregat A, Jupe S, Matthews L, et al: The Reactome Pathway Knowledgebase. Nucleic Acids Res 46:D649–D655, 2018

18. Schaefer CF, Anthony K, Krupa S, et al: PID: the Pathway Interaction Database. Nucleic Acids Res 37:D674–9, 2009

19. Hornbeck PV, Zhang B, Murray B, et al: PhosphoSitePlus, 2014: mutations, PTMs and recalibrations. Nucleic Acids Res 43:D512–20, 2015

20. Romero P, Wagg J, Green ML, et al: Computational prediction of human metabolic pathways from the complete human genome. Genome Biol 6:R2, 2005

21. Mi H, Huang X, Muruganujan A, et al: PANTHER version 11: expanded annotation data from Gene Ontology and Reactome pathways, and data analysis tool enhancements. Nucleic Acids Res 45:D183–D189, 2017

22. Subramanian A, Tamayo P, Mootha VK, et al: Gene set enrichment analysis: a knowledge-based approach for interpreting genome-wide expression profiles. Proc Natl A cad Sci U S A 102:15545–15550, 2005

23. Thiele I, Swainston N, Fleming RMT, et al: A community-driven global reconstruction of human metabolism. Nat Biotechnol 31:419–425, 2013

24. Davis AP, Grondin CJ, Johnson RJ, et al: The Comparative Toxicogenomics Database: update 2017. Nucleic Acids Res 45:D972–D978, 2017

25. Wrzodek C, Büchel F, Ruff M, et al: Precise generation of systems biology models from KEGG pathways. BMC Syst Biol 7:15, 2013

26. Yamamoto S, Sakai N, Nakamura H, et al: INOH: ontology-based highly structured database of signal transduction pathways. Database 2011:bar052, 2011

27. Kandasamy K, Mohan SS, Raju R, et al: NetPath: a public resource of curated signal transduction pathways. Genome Biol 11:R3, 2010

28. Pico AR, Kelder T, van Iersel MP, et al: WikiPathways: pathway editing for the people. PLoS Biol 6:e184, 2008

29. Finn RD, Bateman A, Clements J, et al: Pfam: the protein families database. Nucleic Acids Res 42:D222–30, 2014

30. Carbon S, Mungall C: Gene Ontology Data Archive [Internet], 2018Available from: http://dx.doi.org/10.5281/ZENODO.2529950

31. Hubbard T, Barker D, Birney E, et al: The Ensembl genome database project. Nucleic Acids Res 30:38–41, 2002

32. Rodriguez MA: The Gremlin Graph Traversal Machine and Language [Internet]. arXiv [csDB], 2015Available from: http://arxiv.org/abs/1508.03843

33. McKinney W, Others: Data structures for statistical computing in python, in Proceedings of the 9th Python in Science Conference. Austin, TX, 2010, pp 51–56

34. Messina A, Pribadi H, Stichbury J, et al: BioGrakn: A Knowledge Graph-Based Semantic Database for Biomedical Sciences, in Complex, Intelligent, and Software Intensive Systems. Springer International Publishing, 2018, pp 299–309

35. Messina A, Fiannaca A, La Paglia L, et al: BioGraph: a web application and a graph database for querying and analyzing bioinformatics resources. BMC Syst Biol 12:98, 2018

36. Pareja-Tobes P, Tobes R, Manrique M, et al: Bio4j: a high-performance cloud-enabled graph-based data platform [Internet]. bioRxiv 016758, 2015[cited 2019 Sep 13] Available from: https://www.biorxiv.org/content/10.1101/016758v1

37. Belleau F, Nolin M-A, Tourigny N, et al: Bio2RDF: towards a mashup to build bioinformatics knowledge systems. J Biomed Inform 41:706–716, 2008

38. Himmelstein DS, Lizee A, Hessler C, et al: Systematic integration of biomedical knowledge prioritizes drugs for repurposing [Internet]. Elife 6, 2017 Available from: http://dx.doi.org/10.7554/eLife.26726

39. Flicek P, Amode MR, Barrell D, et al: Ensembl 2011. Nucleic Acids Res 39:D800–6, 2011

40. Ashburner M, Ball CA, Blake JA, et al: Gene ontology: tool for the unification of biology. The Gene Ontology Consortium. Nat Genet 25:25–29, 2000

41. Mungall CJ, McMurry JA, Köhler S, et al: The Monarch Initiative: an integrative data and analytic platform connecting phenotypes to genotypes across species. Nucleic Acids Res 45:D712–D722, 2017

42. Liberzon A, Subramanian A, Pinchback R, et al: Molecular signatures database (MSigDB) 3.0. Bioinformatics 27:1739–1740, 2011

43. El-Gebali S, Mistry J, Bateman A, et al: The Pfam protein families database in 2019. Nucleic Acids Res 47:D427–D432, 2019

44. Kim S, Thiessen PA, Bolton EE, et al: PubChem Substance and Compound databases. Nucleic Acids Res 44:D1202–13, 2016

45. Cotto KC, Wagner AH, Feng Y-Y, et al: DGIdb 3.0: a redesign and expansion of the drug-gene interaction database. Nucleic Acids Res 46:D1068–D1073, 2018

46. Edwards AM, Isserlin R, Bader GD, et al: Too many roads not taken. Nature 470:163–165, 2011

47. Bento AP, Gaulton A, Hersey A, et al: The ChEMBL bioactivity database: an update. Nucleic Acids Res 42:D1083–90, 2014

48. Ainscough BJ, Griffith M, Coffman AC, et al: DoCM: a database of curated mutations in cancer. Nat Methods 13:806–807, 2016

49. Pawson AJ, Sharman JL, Benson HE, et al: The IUPHAR/BPS Guide to PHARMACOLOGY: an expert-driven knowledgebase of drug targets and their ligands. Nucleic Acids Res 42:D1098–106, 2014

50. Simon GR, Somaiah N: A tabulated summary of targeted and biologic therapies for non-small-cell lung cancer. Clin Lung Cancer 15:21–51, 2014

51. Rask-Andersen M, Masuram S, Schiöth HB: The druggable genome: Evaluation of drug targets in clinical trials suggests major shifts in molecular class and indication. Annu Rev Pharmacol Toxicol 54:9–26, 2014

52. Rask-Andersen M, Almén MS, Schiöth HB: Trends in the exploitation of novel drug targets. Nat Rev Drug Discov 10:579–590, 2011

53. Zhu F, Han B, Kumar P, et al: Update of TTD: Therapeutic Target Database. Nucleic Acids Res 38:D787–91, 2010

