## Supplementary material for "Exploring Integrative Analysis using the BioMedical Evidence Graph": IPython Examples: Examples.ipynb

```
# %% [code]
import pandas
import gripql
conn = gripql.Connection("https://bmegio.ohsu.edu/api", credential_file='bmeg_credentials.json')

# %% [code]
G = conn.graph("bmeg_rc3")

# %% [markdown]
## Query 1

# %% [code]
q = G.query().V("Project:TCGA-BRCA").\
out("cases").out("samples").\
has(gripql.eq("gdc_attributes.sample_type", "Primary Tumor")).\
out("aliquots").out("somatic_callsets").outE("alleles").\
has(gripql.contains("methods", "MUTECT")).\
aggregate(gripql.term("geneCount", "ensembl_gene"))

# %% [code]
res = list(q)
df = pandas.DataFrame(res[0].geneCount.buckets).set_index("key")
print(df[:10])

# %% [markdown]
## Query 2

# %% [code]
genes = list(df.index)
q = G.query().V(genes).as_("gene").\
out("pathways").render(["$gene._gid", "$._gid"])

# %% [code]
res = list(q)
df = pandas.DataFrame(res)
print(df[:10])

# %% [markdown]
## Query 3

# %% [code]
q = G.query().V().hasLabel("Pathway").as_("pathway")\
.out("genes").select("pathway")\
.aggregate(gripql.term("pathwayGeneCount", "_gid"))\

# %% [code]
res = q.execute()
df = pandas.DataFrame(res[0].pathwayGeneCount.buckets)
print(df[:10])

# %% [markdown]
## Query 4

# %% [code]
q = G.query().V("Project:TCGA-BRCA").\
out("cases").out("samples").\
has(gripql.eq("gdc_attributes.sample_type", "Primary Tumor")).\
out("aliquots").out("somatic_callsets").out("alleles").\
out("g2p_associations").out("publications").\
aggregate(gripql.term("pub", "_gid"))

# %% [code]
res = q.execute()
df = pandas.DataFrame(res[0].pub.buckets)
print(df[:10])

# %% [markdown]
## Query 5

# %% [code]
q = G.query().V("Project:TCGA-BRCA").\
out("cases").as_("case").out("samples").\
has(gripql.eq("gdc_attributes.sample_type", "Primary Tumor")).\
out("aliquots").out("somatic_callsets").out("alleles").\
out("g2p_associations").out("compounds").\
distinct(["$case._gid", "_gid"]).\
aggregate(gripql.term("compound", "_gid"))

# %% [code]
res = q.execute()
df = pandas.DataFrame(res[0].compound.buckets)
compounds = list(df["key"][:20])
print(compounds)

# %% [markdown]
## Query 6

# %% [code]
res = G.query().V(compounds).as_("compound").out("projects").\
has(gripql.eq("project_id", "CTRP_Breast_Cancer")).\
select("compound").render(["_gid", "synonym"])

# %% [code]
df = pandas.DataFrame(res.execute())
print(df)

# %% [markdown]
## Query 7

# %% [code]
res = G.query().V("Program:CTRP").out("projects")\
.out("cases").out("samples").as_("sample")\
.out("aliquots").out("drug_response").as_("response")\
.out("compounds").hasId("Compound:CID104741")\
.render(["$sample._gid","$response.submitter_compound_id","$response.ec50"])

# %% [code]
df = pandas.DataFrame(res.execute())
print(df[:10])

# %% [code]
samples = list(df[0])

# %% [markdown]
## Query 8

# %% [code]
res = G.query().V(samples).as_("sample").\
out("case").out("same_as").\
out("samples").out("aliquots").out("gene_expressions").as_("exp").\
render( ["$sample._gid", "$exp._data.values"])

# %% [code]
df = pandas.DataFrame(dict(res.execute()))
print(df.iloc[:10,:10])

# %% [code]
```
